# *Clostridium difficile* binary toxin (CDT) is recognized by the TLR2/6 heterodimer to induce an NF-κB response

**DOI:** 10.1101/2020.09.08.288209

**Authors:** Morgan Simpson, Alyse Frisbee, Pankaj Kumar, Carsten Schwan, Klaus Aktories, William A. Petri

## Abstract

*Clostridioides difficile* infection (CDI) represents a significant burden on the health care system, one that is exacerbated by the emergence of binary toxin (CDT)-producing hypervirulent *C. difficile* strains. Previous work from our lab has shown that TLR2 recognizes CDT to induce inflammation. Here we explore the interactions of CDT with TLR2 and the impact on host immunity during CDI. We found that the TLR2/6 heterodimer, not TLR2/1, is responsible for CDT recognition, and that gene pathways including NF-κB and MAPK downstream of TLR2/6 are upregulated in mice with intact TLR2/6 signaling during CDI.

## Background

*Clostridium difficile* is an anaerobic, Gram-positive, spore-forming bacterium that, in the presence of antibiotic-induced dysbiosis, can cause life-threatening colitis. Despite available antibiotic therapy, *C. difficile* is responsible for an estimated 500,000 infections and 13,000 deaths annually in the US [1]. In the past few decades, there has been an emergence of hypervirulent strains thought to be associated with increased disease severity and patient mortality [2]. In addition to expressing the primary clostridial toxins Toxin A and Toxin B, these strains also express a third toxin named binary toxin. This binary toxin consists of an enzymatic component, CDTa, and a binding component, CDTb, which act together to intoxicate intestinal epithelial cells alongside Toxin A and Toxin B. A host receptor for CDT is the lipolysis-stimulated lipoprotein receptor (LSR). Following the heptamerization and association of CDTb to LSR, CDTa binds to the CDTb heptamer and the complex is endocytosed into the cell. Endosomal acidification triggers insertion of CDTb into the endosomal membrane, forming a pore to allow CDTa entry into the host cell cytoplasm, where it inhibits actin polymerization. This ultimately leads to cytoskeletal collapse, cell rounding, and cell death [3].

The intoxication of intestinal epithelial cells by CDT, as well as Toxin A and Toxin B, disrupts the intestinal epithelial barrier, leading to translocation of commensal microbiota, production of inflammatory cytokines and chemokines, and recruitment of inflammatory immune cells to the site of infection. Because of this, the virulence factors produced by *C. difficile* during infection have an important role in host outcome during infection.

Another vital factor is the host immune response, which can be either protective or detrimental to the host [4,5]. Toll-like receptors (TLRs), a class of pattern recognition receptors (PRRs) expressed on the plasma membrane, serve as important frontline responders within the innate immune system, due to their ability to recognize and respond to pathogen-associated molecular patterns (PAMPs), such as bacterial lipoproteins [6]. Previously, our lab has shown that toll-like receptor 2 (TLR2) is capable of recognizing CDT to induce an IL-1β response [7]. However, TLR2 is unique within the TLR family in that it requires heterodimerization with TLR1 or TLR6 in order to initiate a signaling cascade and subsequent downstream immune response [6], and it remains unknown which of these heterodimers is responsible for recognition of CDT.

In this study, we sought to further explore the interaction of TLR2 with CDT, and the potential downstream impact of TLR2 signaling on the host immune response to *C. difficile* infection (CDI). By utilizing a TLR2 reporter cell line along with blocking antibodies against TLR1 and TLR6, we were able to determine that it is the TLR2/6 heterodimer, not TLR2/1, that is capable of recognizing CDT and inducing NF-κB activation. In addition, we used transcriptomic analysis to show that a wide variety of immune-related pathways and genes are upregulated in mice with intact TLR2/6 signaling during infection with a CDT-expressing strain of *C. difficile*.

## Methods

### Cell Culture and Toxins

HEKBlue-hTLR2 reporter cells were obtained from Invivogen and grown in DMEM supplemented with 4.5 g/L glucose, 2 mM L-glutamine, 10% FBS, 100 U/mL penicillin, 100 μg/mL streptomycin, 100 μg/mL Normocin (Invivogen), and 1X HEK-Blue Selection (Invivogen). Cells were rinsed with warm PBS, detached with a cell scraper and resuspended at a density of 2.8 × 10^5^ cells/mL in HEK-Blue Detection media (Invivogen). Neutralizing antibodies against TLR2, TLR1, or TLR6 were added to the cells at a concentration of 5 μg/mL and incubated for 1 hr at 37 °C. Following the incubation, Pam3CSK4 (10 ng/mL) (Invivogen), FSL-1 (10 ng/mL) (Invivogen), or CDTa and CDTb (5 ng/mL each) was added. An equivalent volume of endotoxin-free water (Fisher Scientific) (20 uL) was used as a negative control. Cells were incubated at 37 °C for 6-16 hours. Secreted embryonic alkaline phosphatase (SEAP) was quantified by taking the OD at 655 nm in a spectrophotometer. Purified CDTa and CDTb were expressed in B. megaterium and purified as described previously [8].

### Mice and *Clostridium difficile* Infection

Experiments were carried out using 8- to 12-week-old male and female C57BL/6J mice from the Jackson Laboratory. All animals were housed under specific-pathogen free conditions at the University of Virginia’s animal facility, and all procedures were approved by the Institutional Animal Care and Use Committee at the University of Virginia. Mice were infected using a previously established murine model for CDI [7]. Six days prior to infection, mice were given an antibiotic cocktail within drinking water consisting of 45 mg/L vancomycin (Mylan), 35 mg/L colistin (Sigma), 35 mg/L gentamicin (Sigma), and 215 mg/L metronidazole (Hospira). Three days later, mice were switched to regular drinking water for two days and the day prior to infection, given a single IP injection of 0.016 mg/g clindamycin (Hospira). The day of infection, mice were orally gavaged with 1 × 10^3^ vegetative *C. difficile* (R20291 strain). Mice were euthanized on day 3 post-infection and cecal tissue was harvested for transcriptome analysis.

### Transcriptome Microarray

WT and TLR2^−/−^ mice were infected with the CDT-expressing R20291 *C. difficile* strain (CDT+). The whole-cecal tissue transcriptomic analysis was performed on day 3 post infection. Affymetrix Gene Chips^®^ WT PLUS Regent Kit was used to process the RNA samples. Samples were hybridized to the Affymetrix Mouse Gene 2.0 ST GeneChip^®^. We had six replicates for WT (GEO ID = GSM3452975, GSM3452976, GSM3452977, GSM3452978, GSM3452979 and GSM3452980) and six replicates for TLR2^−/−^ (GEO ID = GSM3452963, GSM3452964, GSM34565, GSM3452966, GSM3452967 and GSM3452968). Differential gene expression analysis was performed on the grouped samples using the limma package in R (with GEO2R functionality on GEO website). Volcano plot was generated using DESeq2 package in R [9]. Genes that had p-value <=0.05 and were 1.5-fold up or downregulated were used for pathway enrichment analysis. Pathway enrichment was done using the ConsensusPathDB database (http://cpdb.molgen.mpg.de) [10].

### Statistical Analysis

Statistical analysis was calculated and significance determined (P<0.05) using ANOVA for multiple comparisons. All statistical analysis was performed using GraphPad Prism software.

## Results

### The TLR2/6 heterodimer recognizes CDT to activate NF-κB

To test if TLR2/1 or TLR2/6 recognizes and responds to CDT, we utilized the HEKBlue-hTLR2 reporter cell line, which has been transfected with human CD14/TLR2 and a secreted embryonic alkaline phosphatase (SEAP) reporter for NF-κB. Stimulation with Pam3CSK4, a TLR2/1 ligand, or FSL-1, a TLR2/6 ligand induced significant NF-κB activation which was reversed following treatment with neutralizing antibodies against the appropriate heterodimer (Fig 1a). Using this system we investigated the TLR2 heterodimer responsible for CDT recognition. Treatment with the anti-TLR2 and anti-TLR6 antibodies led to a significant decrease in NF-κB activation following CDT treatment, while treatment with an anti-TLR1 antibody had no significant effect (Fig 1b). We concluded that the TLR2/6 heterodimer, not TLR2/1, recognized CDT and induced an NF-κB response.

**Figure 1.**
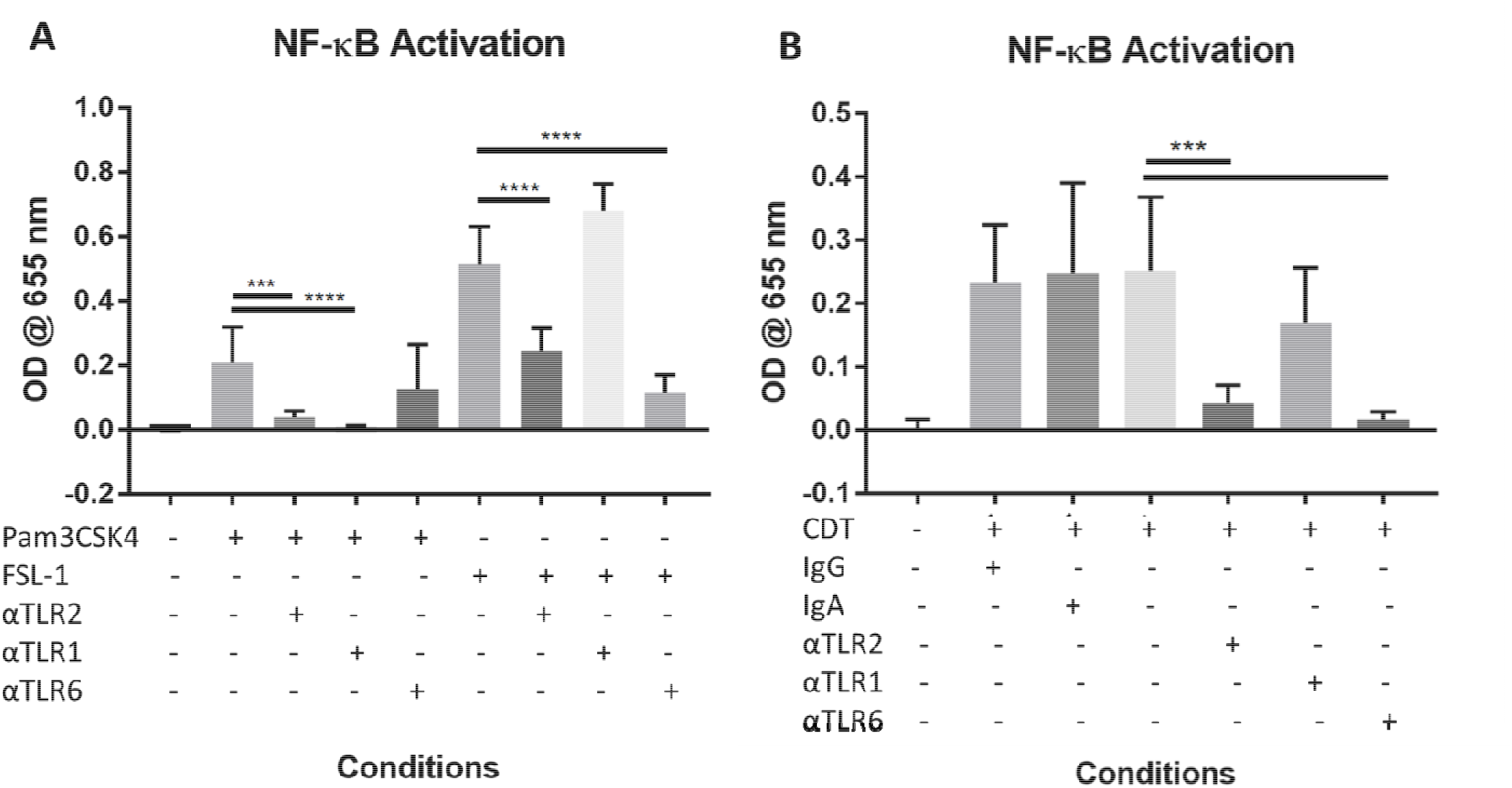
*Clostridium difficile* binary toxin (CDT) induces NF-κB activation via recognition by the TLR2/6, but not the TLR2/1, heterodimer. A: HEKBlue-hTLR2 cells were incubated with 5 μg/mL of anti-TLR2, anti-TLR1, or anti-TLR6 antibodies, treated with 10 ng/mL of Pam3CSK4 or FSL-1, then incubated for 6-16 hours. NF-κB activation was detected by measuring secreted embryonic alkaline phosphatase (SEAP) in the culture media B: HEKBlue-hTLR2 cells were incubated with 5 μg/mL of anti-TLR2, anti-TLR1, or anti-TLR6 antibodies, treated wi h 5 ng/mL of CDTa, CDTb, or full holotoxin, then incubated for 9 hours. NF-κB activation was detected by measuring the SEAP reporter.

### Infection with a CDT-expressing strain of *C. difficile* induces upregulation of immune pathway-related genes

To investigate how TLR2/6 signaling affected downstream gene expression during *C. difficile* infection, we infected WT and TLR2^−/−^ mice with the CDT-expressing epidemic PCR-ribotype 027 strain R20291 and harvested cecal tissue on day 3 post-infection. At this time point, the average weight of the WT mice was 85% of their starting weight, and the average weight of the TLR2^−/−^ mice was 80% of their starting weight (data not shown), indicating that both groups were experiencing severe disease. We compared whole-cecal tissue transcriptomes of WT vs TLR2^−/−^ mice via Affymetrix microarray. Several immune-related genes and pathways were upregulated in WT mice as compared to TLR2^−/−^ mice (Fig 2a-b), demonstrating that the presence of intact TLR2/6 signaling in mice during infection with a CDT-expressing strain of *C. difficile* had a broad impact on the host immune response. Both Cxcl9 and Cxcl10, which encode for chemokines involved in immune cell chemotaxis, were among the upregulated genes. Multiple genes encoding for immunity-related GTPase (IRG) proteins were also upregulated, including Igtp, Iigp1, and Irgm2. One of the genes downregulated in WT mice as compared to TLR2^−/−^ mice was Ly6d, which has been used as a marker for early B cell specification [11].

**Figure 2.**
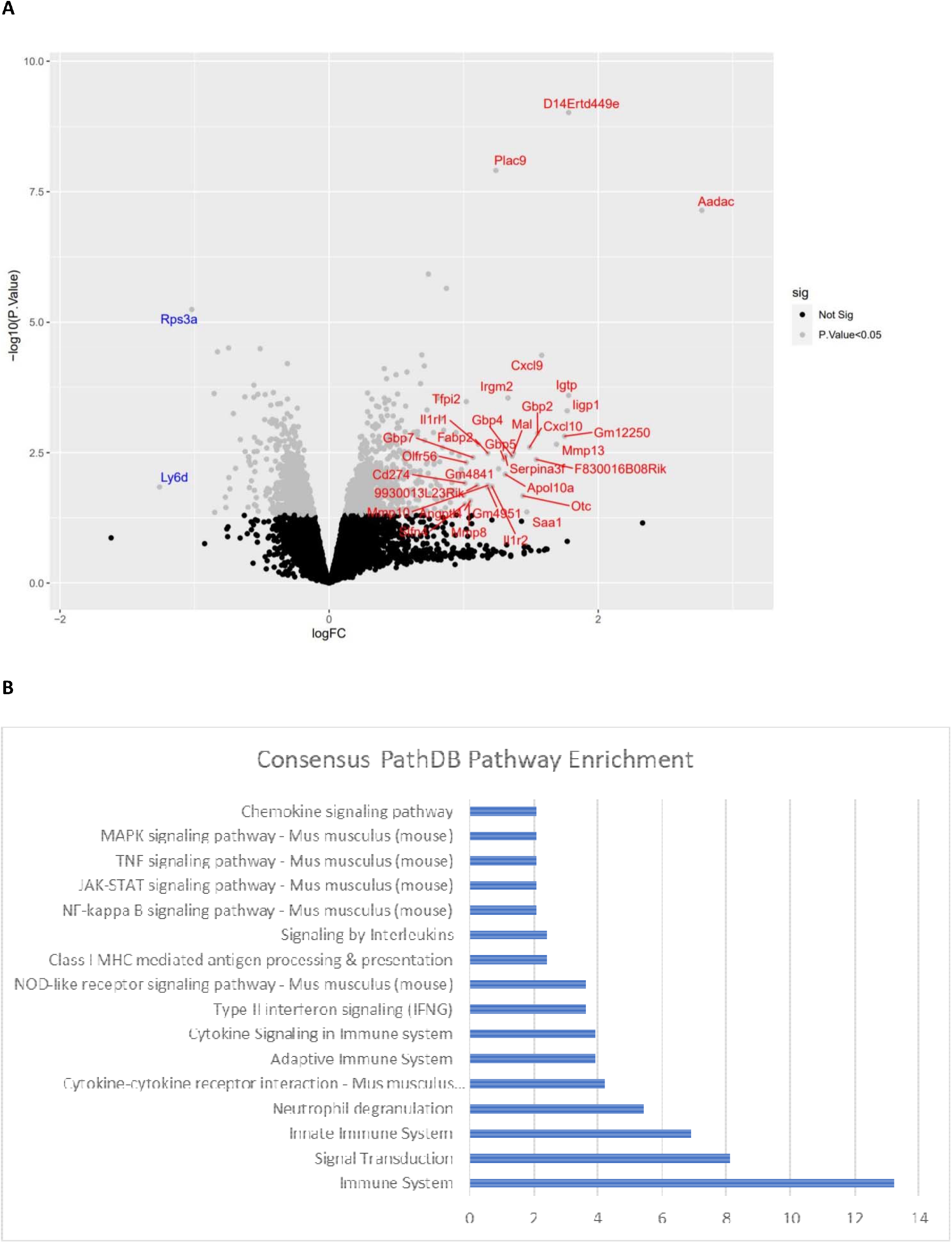
Intact TLR2 signaling during infection with a CDT-expressing strain of C. difficile induces significant upregulation of immune pathway-related genes. WT and TLR2^−/−^ mice were infected with CDT-expressing R20291 and whole-cecal transcriptome analysis was done on day 3 post infection using the Affymetrix Mouse Gene 2.0 ST GeneChip^®^ A: Volcano plot highlighting genes that are upregulated (red) and downregulated (blue) in WT mice compared to TLR2^−/−^ mice infected with CDT-expressing strain of C. difficile (n = 6 animals). On X-axis, logFC is the log2 fold change. Significance of up/down regulation is shown on Y axis (P-value). B: Pathway enrichment analysis using ConsensusPathDB database considering all transcripts with p-value <0.05 and 1.5-fold up/down regulated in WT-CDT1_vs_TLRKO-CDT1. Significance of the identified pathway is shown as P value (−log10P) on the Y axis.

We performed a gene pathway enrichment analysis and found several pathways involved in innate and adaptive immune responses to be enriched in mice with intact TLR2/6 signaling. The enriched pathways included neutrophil degranulation, type II interferon signaling, NOD-like receptor signaling, and class I MHC-mediated antigen processing and presentation. Signaling pathways involving NF-κB, JAK-STAT, PI3k-Akt, TNF, and MAPKs were also upregulated (Fig 1b), indicating that intact TLR2/6 signaling during *C. difficile* infection had an extensive effect on the host immune response.

## Discussion

In this study, we demonstrated that that the TLR2/6 signaling heterodimer recognized CDT to induce NF-κB, and that intact TLR2 signaling in mice had a broad impact on the host immune response during infection with a CDT-expressing strain of *C. difficile*.

We had previously shown that TLR2 can recognize CDT to induce downstream inflammation [6]. TLR2 is unique among the TLRs in that it requires heterodimerization with either TLR1 or TLR6 to initiate downstream signaling cascades and a subsequent proinflammatory response [7]. These heterodimers recognize unique ligands and induce differential downstream immune responses. In this study, we found that blocking either TLR2 or TLR6 with a neutralizing antibody following CDT treatment diminished the downstream inflammatory response, indicating that CDT acts as a TLR2/6 ligand.

Understanding the specifics of TLR2 signaling in the context of *C. difficile* infection can help shed light on the downstream host response, as activation of the TLR2/1 or TLR2/6 heterodimer leads to differing immune responses depending on the context. For example, in a murine model of Y. pestis, TLR2/1 activation has been associated with IL-12p40 producing dendritic cells and inflammatory IFN-γ^+^ T cells, while TLR2/6 activation has been shown to induce IL-10 production and type-1 regulatory T cells [12]. Alternatively, TLR2/6 activation in CD4+ T cells derived from murine lymphoid tissue promoted generation of Th17 cells [13], a cell type our lab has shown to be detrimental to the host during CDI [5]. It not yet known how precisely stimulation of TLR2/1 or TLR2/6 can lead to such varying downstream immune responses in different contexts, however, it is clear that understanding which TLR2 heterodimer is involved in the immune response in a specific context can aid in the development of targeted immunotherapies.

To further explore the impact of TLR2/6 signaling on the host immune response during CDI, we performed a transcriptome array of WT and TLR2^−/−^ mice during infection with a CDT-expressing strain of *C. difficile*. Among the gene pathways upregulated in mice with intact TLR2/6 signaling were MAPK and NF-κB signaling, neutrophil degranulation, cytokine signaling, NOD-like receptor signaling, type II interferon signaling, class I MHC-mediated antigen processing and presentation, chemokine signaling, and TNF signaling. This result highlighted the potential contribution that TLR2/6 signaling has on the inflammation seen during CDI, and as such, may be a potential target for therapeutic intervention.

While this work has furthered our understanding regarding the influence of CDT on TLR2/6 signaling and its impact on immunity-related gene expression, there are still some avenues to be investigated. For example, we have demonstrated that CDT interacts with TLR2/6 to instigate a downstream response, but it is unclear whether this interaction is direct or indirect through as yet unknown mediators. It is known that CDT binds to LSR [14], however, how or if LSR plays a role in the interaction between CDT and TLR2 has yet to be investigated. Finally, it remains unclear whether the interaction of CDT with LSR takes place at the cell surface, or within the endosome. These questions all represent possible areas of future study.

Overall, we have demonstrated that the TLR2/6 heterodimer is able to recognize and respond to CDT in vitro to induce inflammation, and that intact TLR2 signaling in mice during infection with a CDT-expressing strain of *C. difficile* led to upregulation of a variety of immune-related gene pathways. Understanding this interaction and how it may impact downstream immune responses may aid in the design of more targeted therapeutics to dampen damaging inflammation during infection.

## Acknowledgements

The authors would like to acknowledge Carrie Cowardin, Mahmoud Saleh and Alexandra Donlan for scientific advice and discussion.

## Meeting(s) where manuscript was previously presented

International Congress of Mucosal Immunology, July 2019, Brisbane, Australia

Mucosal Immunology Course and Symposium, July 2018, Oxford, England

